# Sex-bias in utero alters ovarian reserve but not uterine capacity in female offspring

**DOI:** 10.1101/2022.07.11.499550

**Authors:** Annika V Geijer-Simpson, Haidee Tinning, Tiago H C de Bem, Ioannis Tsagakis, Alysha S Taylor, Laura Hume, Lisa M Collins, Niamh Forde

## Abstract

Environmental stressors to which a foetus is exposed, affect a range of physiological functions in post-natal offspring. Such stressors include disproportionate steroid hormone concentrations in the uterine environment. We aimed to determine the *in-utero* effect of steroid hormones on reproductive potential of female offspring using a porcine model. Hypothesising that an *in-utero* sex bias will influence ovarian reserve and endometrial morphology in the breeding gilt. Reproductive tracts of pigs from female-biased litters (>65% female, n=15), non-biased litters (45-54.9% female, n=15), and male-biased litters (<35% females, n=9) were collected at slaughter (95-115 kg). Ovaries and uterine horns were processed for histological approaches and stained using H&E or IHC techniques. All measurements were conducted in QuPath (Bankhead et al, 2017). Variability of data within groups was analysed with a Levenes test, whilst data was analysed using linear models in R. In the ovarian reserve, there was a significant interaction between the birth weight and the sex ratio of a litter from which a pig originated (p=.015), with low-birth-weight pigs from male-biased litters having a higher number of primordial follicles and the opposite trend seen in pigs from female-biased litters. This was not reflected in recruited, nor atretic follicles. In the uterine horn sex bias held no effect on development as seen in this study. Birth weight held more effects on the gilts. A lower BW decreased the proportion of glands found in the endometrium (p=.045). BW was found to be far more variable in both male-biased and female-biased litters (p=.026). The variability of primordial follicles from male-biased litters was greater than non-and female-biased litters (p=.014). Similarly, endometrial stromal nuclei had a greater range in male- and female-biased litters than non-biased litters (p=.028). There was a greater effect on both ovarian reserve and uterine development of piglet BW than the litter bias. There seems a benefit of being androgenised on ovarian reserve whilst no effects were found for the morphology or endometrial gland proliferation of the uterine horns. However, a crucial finding was in the variability of the data. Both primordial follicles in the male-biased ovary, and stromal nuclei in the male- and female-biased uterine horns had a wider spread in numbers than non-biased litters. This could be inflating the variability of reproductive success seen in animals form male-biased litters by two means. Firstly, by a higher likelihood of insufficient primordial pools. Secondly, through a potential impact on stromal-derived growth factors or insufficient support of the underlying implantation structures, leading to an increased variability in uterine implantation capabilities, and thus survival of the embryo.

## INTRODUCTION

The Developmental Origins of Health and Disease (DOHaD) describes how the developmental plasticity and *in-utero* programming of offspring could contribute to susceptibility of a range of adult disease with the maternal *intra-uterine* hormonal environment has been shown to influence many aspects of offspring development, some of which are sexually dimorphic in nature [1]. Studies specifically investigating the influence of foetal hormones have indicated effects such as endocrine alterations [2], altered physiological development such as delayed lung maturation [3], behavioural changes such as increased aggression [4], and reduced maternal and paternal behaviour [5]. These effects are observed following an androgenisation of the uterine environment in litter bearing species, often caused by disproportionate numbers of males *in-utero*. This may be a consequence of either; females positioned between males (which is proportionately more likely to happen in male-biased litters) or; the overall proportion of males excreting testosterone from the point of sexual differentiation [6]. A biased litter, one that skews toward a predominantly androgenised or oestrogenised environment (those consisting of >60% of one sex), occur frequently both within commercial production systems and among wild pig populations, where proportions of 1.3:1 males to females per litter have been reported [7]. In the commercial pig a sex-biased *in utero* environment hormonal environment has been shown to affect several aspects of reproductive function in the offspring. Specifically, females that originate from male-biased litters have fewer teats [8], a lower conception rate at first mating [9], increased sensitivity to gonadotropins [10], and altered LH surge profiles [2], compared to offspring originating from female-biased litters. The cause of these changes in reproductive efficacy remains unclear.

Arguably, the most vital organisational event in female reproductive development during gestation is the developing Primordial Germ Cells (PGC). PGCs are originally derived from the proximal epiblast cells of pre-gastrulating embryos [11], i.e. prior to generation of the three primary germ layers. In pigs these have been identified in the dorsal mesentery at embryonic day (ED)18-20 which then migrate to colonise forming the genital ridge at ED23-24 [12]. It is established that the PGC’s undergo epigenetic reprogramming over a period of several weeks, beginning at ED12 [13]. In the pig, meiosis, initiation, and formation of primordial follicles begins at E48 and continues until 25 days post parturition [14]. Differentiation of the Wolffian duct in the male begins at ED26, at which point secretion of testosterone begins [15]. Hence, laying down and formation of the primordial follicles in female offspring occurs once testosterone from male littermates is present in the uterine environment. The point at which the number of PGCs peaks is determined by morphogenesis and germ cell dynamics, and it is still not understood how the ovarian reserve is maintained and what triggers activation of folliculogenesis occurs [16]. It is clear in the pig that there is an effect on ovulation patterns. However, it is currently unclear whether an androgenised uterine environment affects PGC formation or follicular recruitment of the PGCs in the pig.

Data from other species indicates that an androgenised uterine environment effects the development PGC establishment and subsequent follicular recruitment, but also gross morphology of reproductive tracts. The ovulation rate of an individual does impose a clear and direct limitation on potential litter sizes in the pig. However, despite ovulation rates having improved in pigs via selective pressure this hasn’t been reflected in any significant increases to litter sizes [17]. With ovulation and conception rates in the breeding sow being greater than 95% [18], the major limiting factor to maximized litter sizes is embryonic death. Only 30-50% of fertilised ova survive through gestation [17]. This loss is predominantly seen during the pre- and peri-implantation period defined as ED12-18 [17]–[19]. This coincides with conceptus elongation, synthesis and release of oestrogen for maternal pregnancy recognition, and trophectoderm differentiation, which is followed by foetal and epithelial attachment [20]. There is then a secondary wave of embryonic loss at ED 30-40 due to crowded *in-utero* conditions [21]. It is therefore clear that litter sizes, and conceptus survival, is contingent on uterine capacity. This is determined by three main factors; uterine length, uterine blood flow, and uterine gland development [21]. The uterine capacity of a pig is critical it as it contributes to the reproductive potential of the female i.e. even if there are large numbers of developmentally competent embryos present there is a physical limit on the numbers that can implant. It is vital that the uterine tissues can recognise and respond to maternal and conceptus signals crucial for a successful and established pregnancy [21]–[23]. Not only is this vital for pregnancy recognition, but the uterine capacity will define the environment in which foetal development occurs [22]. The functional layer of the uterus that is especially crucial for successful pregnancy outcomes, the endometrium, may be influenced by an androgenised litter. The endometrium is a heterogenous tissue comprised of several different cell types including various secretory cells such as the luminal and glandular epithelium [19]. The endometrial surface of the pig is folded, which at conceptus attachment (ED14) has a conceptus localised increase in endometrial surface folding [24]. The opposite folds interlock, reducing the luminal space throughout the pregnancy, maximising luminal epithelial and foetal contact [19], this is crucial to facilitate maternal and foetal communication. Endometrial glands along with the luminal epithelium secret uterine luminal fluid, a complex array of proteins and related substances [25]. The uterine luminal fluid is critical for endometrial function and conceptus survival as they contain enzymes, growth factors, cytokines, nutrients, transport proteins, and other regulatory molecules (See review by Bazer and Johnson 2014 – “Pig blastocyst-uterine interactions”) [26]. Maternal endometrial gland hyperplasia and hypertrophy is extensive during gestation [27], [28] with large amounts of granular, acid phosphate-positive material within the glands, indicative of a high level of secretory activity [23]. Sheep with blocked uterine horn gland development (UGKO – Sheep Uterine Gland Knock Out) indicate a failure of conceptus elongation at ED14 and are rendered infertile (11). Evidencing the fact that uterine glands are critical mediators for the uterine ability to support a successful pregnancy. Formation of the uterine glands occur in the neonatal piglet by branching and budding of the luminal epithelium [23], reaching histoarchitectural maturity by 120 days of age [27]. Although the glands form neonatally, the histogenesis of the initial uterine horn development, and luminal epithelium, takes place *in-utero* [23], [30] making this process vulnerable to environmental effects of which the foetus is exposed to. The understanding of the impact disruptions of uterine development may hold on endometrial structure and function in adult mammals is crucial for unravelling the high rates of peri-implantation embryonic loss in both livestock and humans [23], as this may render the uterus unable to support especially small for gestational age individuals. For example, adult cows exposed to progesterone and oestradiol benzoate *in-utero* demonstrate a reduced number of endometrial glands [31]–[33]. Furthermore, in pigs, neonatal progesterone treatment initially accelerated gland development, but reduced adult glandular development [34]. Impairment, as described above, has been indicative of a reduced fertile capacity [30].

This study aims to investigate how a sex biased *in-utero* environment may influence the development of both the follicular pool, and endometrial glands; both critical components for successful pregnancy. To do this we investigated the influence of *in-utero* sex ratio a female was gestated in on the (i) follicular patterns in the pig, (ii) uterine morphology and endometrial gland proliferation. This was achieved through investigation of the two following objectives: (i) Investigation of how the established primordial germ cell pool is affected by different *in-utero* hormonal biases and identification of any differences in follicular recruitment, or follicular atresia profiles dependent on litter sex bias, (ii) effects the development of gross uterine structures and level of proliferation within the uterus.

## MATERIAL AND METHODS

After consultation with the University of Leeds Animal Welfare and Ethical Review Committee no ethical approval was sought. It was deemed that due to no interventions taking place, and the pigs remaining fully under the commercial management, ethics was not necessary. This complies with the 3Rs strategy of utilising animals which are already part of a process.

### ANIMAL MODEL

Our model recruited individual female offspring at birth that were retrospectively assigned to one of three experimental groups. Females from 1) female-biased *in utero* litter (>65% females), 2) non-biased (45.9-55% females), or 3) male-biased (<35% females) litters. Only litters with 10 or more live-born piglets were included in the study and mummified piglets were not included as accurate identification of sex was unreliable.

All animals were reared in a closed indoor system at the National Pig Centre on partially slatted floor systems. The pigs used were either not part of any other trials or used as control pigs in other dietary studies. These female pigs (JSR Large White x Landrace females JT dam- line x JSR Pietrain-based Geneconverter 900 sire-line) were tracked through the production system using RFID ear-tags. The pigs were weighed weekly in the month leading up to slaughter in order to optimise an accurate estimate of slaughter weight. At commercial slaughter weight (95-115 kg) pigs were delivered to the abattoir weekly over three collection time points per batch, a total of 10 time points.

### TISSUE COLLECTION AND DISSECTION

To control for within-litter variability, two females were selected at random (using randomizer.org) for reproductive tract collection at slaughter from each litter. Reproductive tract collections occurred over three periods in November 2018, June-August 2019, and July-August 2020. The tracts were collected on the abattoir line and transported to the laboratory on ice within 1.5 hours for tissue processing. The ovaries were dissected from the reproductive tracts and individually weighed. The right ovary was divided in half and placed in 10% neutral buffered formalin. The uterine horns were dissected away from the connective tissue and whole cross sections from 1/3 of the way down from the tip of the uterine horn (UH; Figure 1a) were also fixed in formalin for forty-eight hours. Ovaries and UH were removed from formalin and ovaries further divided in half lengthways through the cortex, resulting in a quartered ovary (**Error! Reference source not found.**b). UH were and ovary were subsequently dehydrated through a series of ethanol washes and imbedded in paraffin for histological examination. In total the reproductive tracts from 39 piglets were assessed; male-biased = 9, non-biased = 15, and female-biased = 15. Unless otherwise stated all chemical were obtained from Thermo Fisher Scientific, Waltham, US.

**Figure 1.**
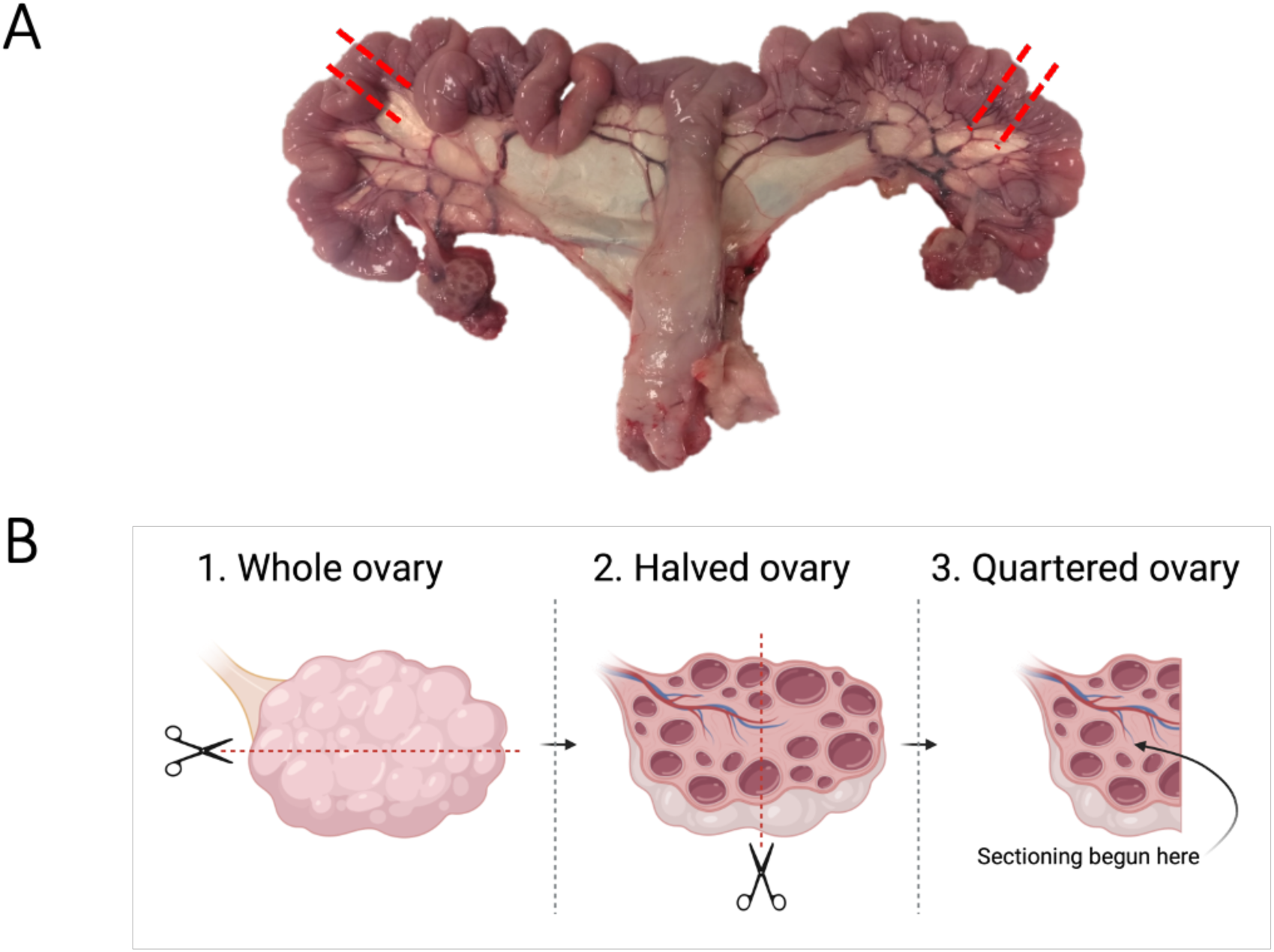
Schematic of how the reproductive tracts were dissected. A) uterine horn sections (1cm) were taken at the marked points, and B) ovaries were dissected, resulting in the final ovary sections. The place at which the first section was taken is marked out under 3. Quartered ovary. Created with Biorender.com.

**Figure 2.**
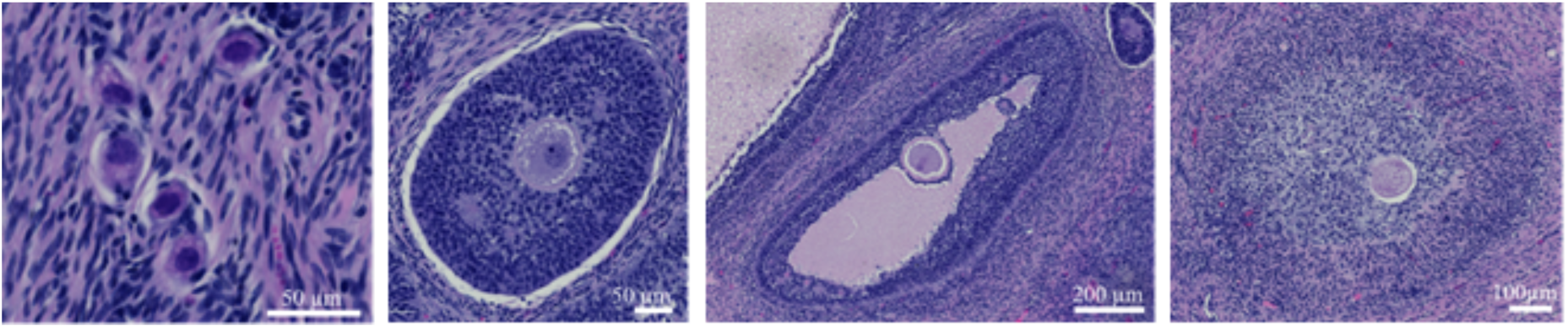
Images taken of follicles representing the different stages of development. A) A cluster of primordial follicles. B) Pre-antral follicle. C) Antral follicle. D) Atretic follicle. Arrowheads indicate location of the oocyte.

### HISTOMORPHOMETRY

#### HEMATOXYLIN AND EOSIN STAIN

For histomorphometry of the ovary, 8 µm serial sections of the embedded ovary from an individual were mounted on polysine-coated Microslides (VWR International, Radnor, US). For UH two individual slides containing whole cross sections of UH per animal were assessed. Slides were de-paraffinised three times in Histo-clear for 10 min. Slides were then hydrated decreasing concentrations of ethanol (3 x 100% EtOH, 2 x 95% EtOH, and once in 70% EtOH). All slides were finally rinsed in three times in tap water. Slides were incubated in hematoxylin (Sigma Aldrich, St. Louis, US - 7g/L) for 5 min, slides rinsed in 3 x in tap water, dipped in 0.25% acid alcohol for 5 sec and immersed in cold water. At this stage stain intensity was checked under a microscope before proceeding. Stain was blued in hot tap water for 1 minute and then rinsed 3 x in cold tap water. Slides were then placed in a Mordant of 95% EtOH. Slides were then stained in Eosin for 10 min, and dehydrated through 2 changes of 95% EtOH, and 3 changes of 100% EtOH. Finally, slides were places in Histo-clear for 10 min x 3. Ten serial sections per ovary per animal, (each 160 µm apart), were selected with the initial section being closest to the ovarian cortex, (**Error! Reference source not found.**). As the distance between sections a full cross section of the oocyte, <110 µm [35], could not be fully present in two analysed sections, ensuring that no included follicle would be counted twice. For the one section per UH of each individual were also stained.

#### IMMUNOHISTOCHEMISTRY – PROLIFERATING CELL NUCLEAR ANTIGEN

For investigating uterine gland proliferation one section per uterine horn were stained using an immunohistochemistry (IHC) technique using the Vector Lab VectaStain Elite ABC-HRP kit. Overnight incubations (4°C) of the sections were carried out with mouse monoclonal anti-proliferating cell nuclear antigen (PCNA) (1:200, Invitrogen). Control sections were incubated with a mouse IgG Isotype Control (1:200, Invitrogen). The sections were then incubated with the secondary antibody and ABC reagent previously described for 60 min in a humidity chamber. Development of sections was carried out using diaminobenzidine substrate (ThermoFisher).

### IMAGE ANALYSES

#### OVARIAN RESERVE

To assess the ovarian reserve the number of primordial, pre-antral, antral and atretic follicles were counted and classified according to Almeida and colleagues [36] (Figure 1). Primordial follicles (Figure 1– A) were identified as an intact oocyte surrounded by a single layer of squamous (pre) granulosa cells. An enlarged oocyte that is surrounded by a single, or multiple, layers of cuboidal granulosa cells was classified as pre-antral (Figure 1-B). Once the follicle had developed a clear antral cavity that was the same size as, or larger than, the oocyte it was classified as an antral follicle (Figure 1– C). The antral follicles have several layers of granulosa cells and have a well-developed thecal layer. All the above follicle types had intact oocytes with no signs of apoptosis or degradation as described by Almeida and colleagues [36]/ If degenerative changes had occurred including reduction of the oocyte or condensation of the nuclear chromatin, or changes to the antral cavity such as scattered granulosa cells, the follicle was identified as atretic (Figure 1-D).

#### GROSS UTERINE MORPHOLOGY AND CELL PROLIFERATION

To assess endometrial capacity, we measured gross morphological structures. As a proxy measure of uterine support we analysed the ratio of secretory cells (luminal and glandular epithelia) to stromal cell area. The surface area of the endometriumper section was measured (cm^2^) along with the perimeter of the luminal gap (cm^2^). Manual counts were made of the glands in the endometrium of total sections, with the number of larger glands also counted. An automated cell nuclei detection within QuPath was used to count the stromal cells within an area of 20,000μm^2^. These structures are visualised in Figure 3. Proliferative capacity of the endometrium was measured using the PCNA IHC stain. An automated DAB stain analysis was used for the entire sections of the uterine horns to measure stain intensity. This allowed for identification of cells with a negative or positive stain, along with division of positively stained cells into a low, moderate, or high stain intensity. Parameters used for the detection were refined using three sections previously stained in the stain optimisation process. The parameters used for stain detection were as follows; pixel size, 0.5µm with thresholds being Low; 0.1-0.3, Moderate; 0.3-0.5, High; 0.8-1.

**Figure 3.**
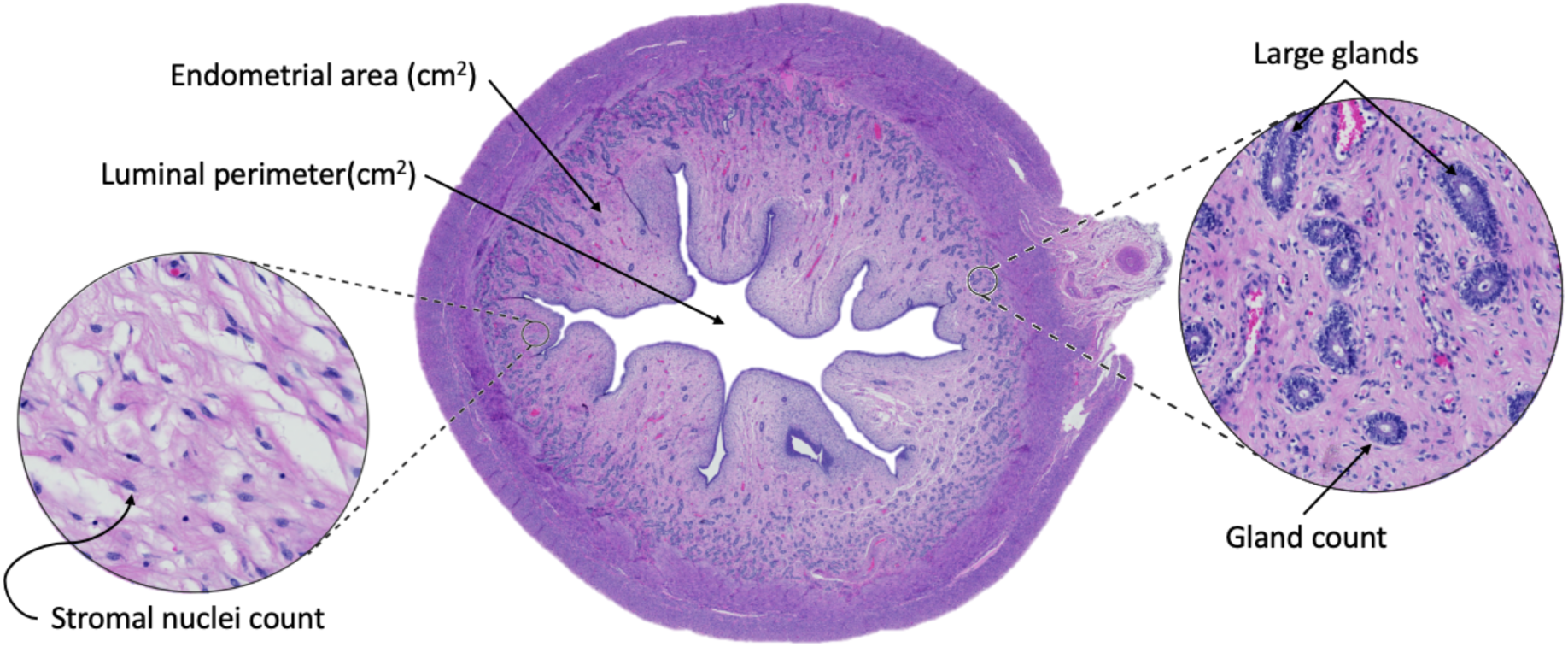
Images taken of uterine horn cross-sections. Example of uterine cross section with an H&E stain and indicators of the morphological structures investigated.

### DATA PROCESSING AND ANALYSIS

All statistical analyses were performed in RStudio [37] using *lme4* [38], and data was plotted using PRISM (GraphPad Prism version 9.0.0 for MacOS, GraphPad Software, San Diego, California USA, www.graphpad.com). Normality was assessed using appropriate tests as detailed below, alongside histograms, and QQ-plots. Data considered to fit a Gamma distribution were tested using **“gamma_test”** in package “**goft**”. Collinearity between predictor values was checked using the “**vif**” function in R package “**car**”. Ovary weight (g) and slaughter age (days) was removed due to collinearity (>3). Critical alpha level was applied as p=0.05. Akaike Information Criterion (AIC) model selection was used to distinguish between a set of possible models, each describing the relationship between the predictor variables. The AIC of the final model used for each analyses is detailed in table 1 and 2. Transformations were required for certain variables to fit their distributions. This was with either a square root transformation, function “sqrt”, or a log transformation, function “log” and is detailed in table 1 and 2.

**Table 1.** The response variables analysed for the ovary and the specific model, model type, distribution of the variable, and AIC value of the most fitting model. All analyses were performed in RStudio and were carried out using lme4. Follicle numbers manually counted on H&E stained histological sections and reproductive tracts were collected from pigs originating from either female biased (n = 15), non biased (n = 15), or male biased groups (n = 9).

| Reponse variable | Model | Model Type | Distribution | AIC value |
| --- | --- | --- | --- | --- |
| Primordial follicles | 1 | GLM | Gamma | 369.0617 |
| Recruited follicles | 1 | GLMM | Gamma | 186.1362 |
| Atretic follicles | 2 | GLM | Gamma | 190.557 |
| Total follicles | 1 | GLM | Gamma | 371.1647 |

The models used were one of the following (specified per response variable in tables 1 and 2);

Models 1- *<Response variable>* per cm^2^ with predictor variables being *litter sex ratio* and *birth weight* as multiplicative, *slaughter weight* as additive, and *litter (ovary) or animal ID (UH)* as random effect.

Models 2 - *<Response variable>* per cm^2^ with predictor variables *litter* as random effect, *litter sex ratio, slaughter weight* as additive, and *litter (ovary) or animal ID (UH)* as random effect.

To investigate whether there was more variability within our investigated response variables between pigs of different *in utero* sex ratios, variance of data points was measured using **“leveneTest”** package.

#### OVARIAN RESERVE

Data was analysed in two ways. Firstly, looking at follicle numbers in the ovary as a whole; and secondly controlling for variations in the manual dissection of the ovary by investigating the number of follicles per cm^2^ of observed tissue. As results were reflective of each other, we have only reported the results per cm^2^ for reader ease.

Gamma regression models were used to test the effect of predictor values on the following response variables; (I) primordial follicle count, (II) recruited follicle count, (III) atretic follicle count, and (IV) total follicular count. All predictor and response variables are described in Table 1.

#### GROSS UTERINE MORPHOLOGY AND PROLIFERATION

Analyses of data was performed to investigate difference of <u>rudimentary morphology</u>. <u>Proportional analyses</u> were conducted to investigate whether there was a difference in the proportion of secretory structures between bias litters. The following comparisons were made; luminal perimeter, stromal cells and endometrial glands in relation to endometrial area. It was then important to compare whether the proportion of these secretory cells differed in proportion to each other between individuals from different *in-utero* sex ratios. These proportional comparisons are all detailed in Table 2. The IHC stained cells were analysed in the same manner as for the morphological analyses. However, low stain intensity was found to hold a normal distribution and was analysed using model 2, and high stain intensity required a square root transformation. This is detailed in table 2.

**Table 2.** The response variables analysed for the uterine horn and the specific model, model type, distribution of the variable, and AIC value of the most fitting model. All analyses were performed in RStudio and were carried out using lme4. Reproductive tracts were collected from pigs originating from either female biased (n = 15), non biased (n = 15), or male biased groups (n = 9). All observations were made from histological sections using either H&E stains or an IHC stain for cell proliferation (PCNA).

| Response variable | Model | Model Type | Distribution | Transformation | AIC value |
| --- | --- | --- | --- | --- | --- |
| Rudimentary analysis |  |  |  |  |  |
| Endometrial area (EA) | 1 | GLM | Gamma |  | -127.258 |
| Luminal perimeter (LP) | 1 | GLM | Gamma | Log | -12.6513 |
| Stromal nuclei (SN) | 2 | GLM | Poisson |  | 575.5295 |
| Gland count (GC) | 2 | GLM | Poisson |  | 1398.412 |
| Large gland count (LGC) | 2 | GLM | Poisson |  | 789.2035 |
| IHC analysis |  |  |  |  |  |
| Positively stained cells | 1 | GLM | Gamma |  | -70.7763 |
| Low intensity | 1 | GLMM | Gaussian |  | -121.477 |
| Moderate intensity | 1 | GLM | Gamma |  | -218.274 |
| High intensity | 1 | GLM | Gamma | Square root | -149.745 |
| Proportional analysis |  |  |  |  |  |
| **LP: EA** | 2 | GLM | Gamma |  | 1476.293 |
| **SN: EA** | 2 | GLM | Gamma |  | 531.1063 |
| **GC: EA** | 2 | GLM | Gamma |  | 1013.363 |
| **SN: LP** | 2 | GLM | Gamma | Square root | 117.1338 |
| **GC: LP** | 2 | GLM | Gamma |  | 749.575 |
| **GC:SN** | 2 | GLM | Gamma |  | 749.575 |

## RESULTS

In total 39 pigs were used for analysis, 15 females from female-biased litters, 15 from non-biased litters, and 9 from male-biased litters. Pigs were excluded from the trial if they failed to reach commercial slaughter, one pig was excluded once reproductive tracts were collected due to an active infection of the tract. The mean birth weight and slaughter weight (± SD) of the pigs were: 1) female-biased - 1.65 kg (± 0.446) and 107 (± 8.140) kg, 2) non-biased - 1.47 (± 0.207) and 110 (± 5.275) kg, and 3) male-biased - 1.51 (± 0.303) and 113 (± 5.915) kg. No significant difference was observed.

The ovarian tissue analysed was on average 16.6 (± 3.197) cm^2^ in female-biased pigs, 17.8 (± 4.386) cm^2^ in non-biased pigs, and 15.0 (± 3.198) cm^2^ in male-biased pigs.

### VARIABILITY OF REPRODUCTIVE PARAMETERS IN INDIVIDUALS

The variability of data points between the sex ratio groups were analysed as visualised in Figure 5. This revealed a bigger range in birth weights (F-value=4.073, df=35, p=.026) of females from both female- and male-biased litters compared to non-biased litters. There was a significantly larger variance of both primordial (F-value=4.801, df=36, p=.014) and total (F-value=5.381, df=36, p=.009) follicle numbers in females from male-biased compared to those that originated from non- or female-biased litters. This also revealed that there was an increased variability in the number of stromal cells in endometria recovered from females from sex-biased litters compared to non-biased (F-value=3.8134, df=71, p=.027). The variability did not differ between litter sex bias in endometrial area (F-value=.2472, df=71, p=.7817), luminal perimeter (F-value=2.3527, df=71, p=.1025), nor total gland (F-value=2.4973, df=71, p=.08951) or large endometrial gland counts (F-value=2.0419, df=71, p=.1373).

**Figure 4.**
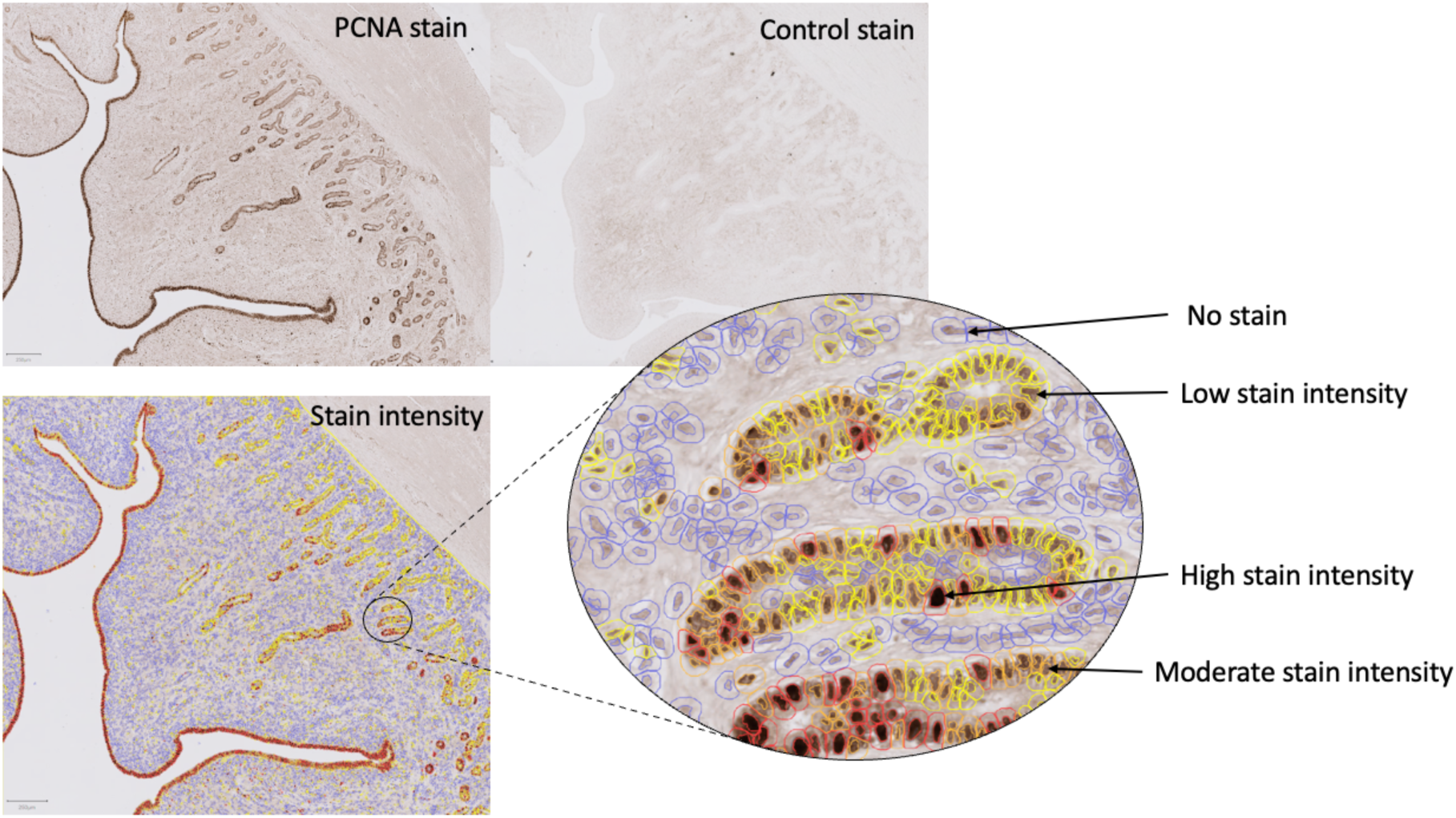
Images of IHC stained sections (8um) using a proliferating cell nuclear antigen antibody, including the identified stain intensity and the control section. The automated stain detection output can be seen including visual representations of cells considered to have no stain, low stain, moderate stain, and high stain intensities.

**Figure 5.**
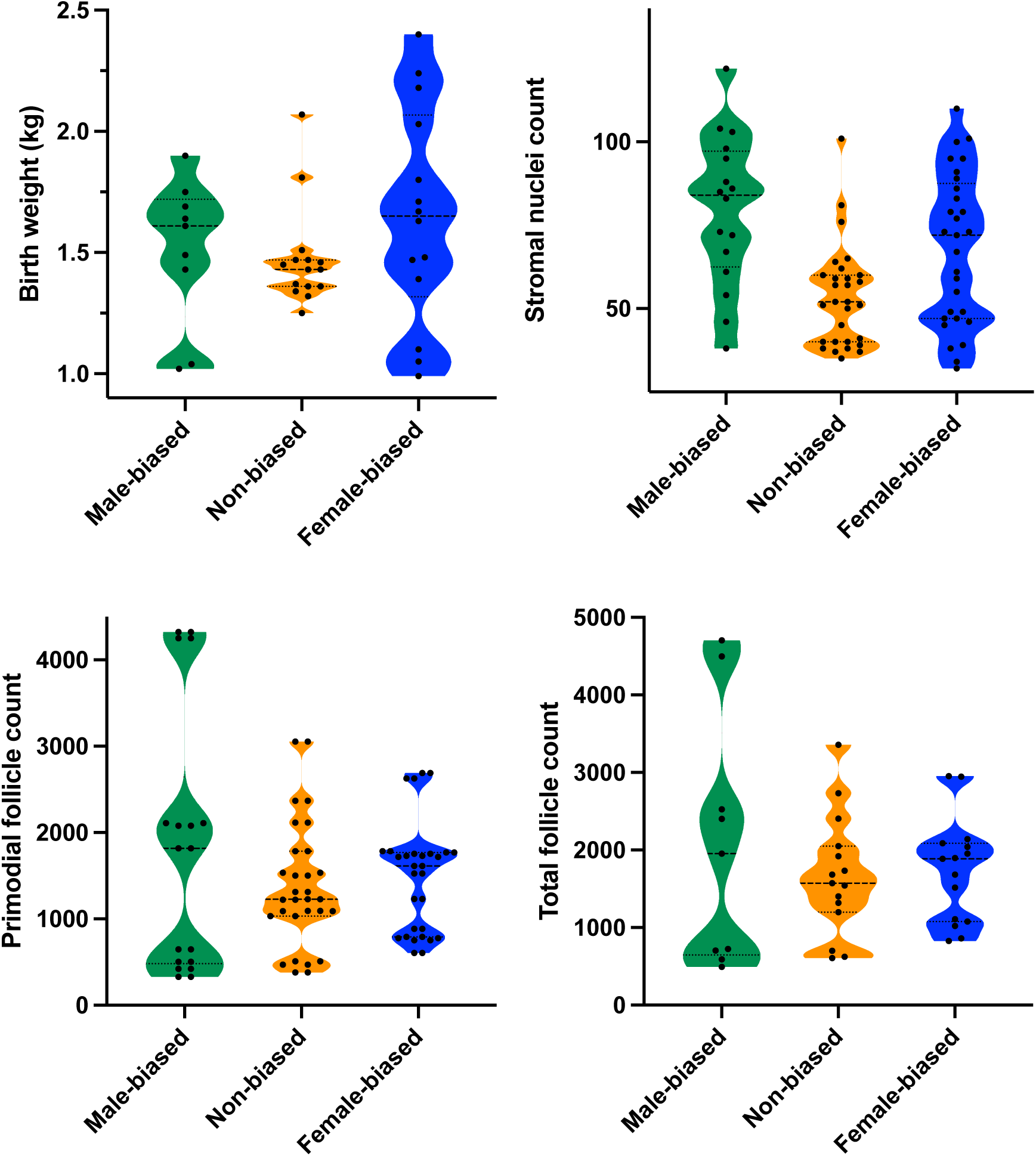
Weight and reproductive parameters from females gestated in different in utero environments. Variability of data represented by a violin plot of the birth weight (kg), stromal nuclei count, primordial follicles, and total follicles between male-biased (n=9), non-biased (n=15), and female-biased (n=15) groups. Data was analysed using a Levenes test. Birth weights of piglets were measured on their first day post parturition post their first suckling event. Stromal nuclei count, primordial follicle counts and total follicle counts were all analysed on H&E stained sections. Manual counts were made for follicles, and an automated nuclei count used in QuPath 0.2.0.

### INTERACTION OF BIRTHWEIGHT AND SLAUGHTER WEIGHT

The following results from the mixed models is summarised in Table 3.

**Table 3.** Mixed model outputs for each response variable. Outputs from the models run for each response variable including the number of animals in the analyses, specific test statistic, and p-value. Signif. Codes: 0 ‘***’; 0.001 ‘**’; 0.01 ‘*’; 0.05 ‘.’.

| Response variable | N of animals / df | Interaction |  | Litter sex bias |  | Birth weight |  | Slaughter weight |  |
| --- | --- | --- | --- | --- | --- | --- | --- | --- | --- |
|  |  | Test statistic | p-value | Test statistic | p-value | Test statistic | p-value | Test statistic | p-value |
| Primordial follicles | 34 | 2.637 | **0.008\*** | NA | NA | NA | NA | 0.541 | 0.589 |
| Recruited follicles | 34 | 0.789 | 0.436 | 0.442 | 0.679 | -0.669 | 0.509 | 0.995 | 0.330 |
| Atretic follicles | 34 | NA | NA | 0.817 | 0.414 | -1.775 | 0.076 | 1.468 | 0.142 |
| Total follicles | 34 | -2.754 | **0.006\*** | NA | NA | NA | NA | -2.754 | 0.611 |
| Endometrial area (EA) | 64 | -1.416 | 0.157 | 0.022 | 0.825 | -1.376 | 0.169 | 0.787 | 0.431 |
| Luminal perimeter (LP) | 64 | -1.595 | 0.1107 | 1.096 | 0.2731 | -1.915 | 0.055 | -0.151 | 0.880 |
| Stromal nuclei (SN) | 64 | NA | NA | -0.287 | 0.774 | -0.916 | 0.359 | 1.237 | 0.216 |
| Gland count (GC) | 64 | NA | NA | 0.117 | 0.907 | 0.058 | 0.954 | 0.54 | 0.957 |
| **Large gland count (LGC)** | 64 | NA | NA | -1.033 | 0302 | 0.432 | 0.666 | -0.908 | 0.364 |
| **LP:EA** | 64 | NA | NA | 0.874 | 0.382 | -0.142 | 0.887 | -1.397 | 0.162 |
| **SN:EA** | 64 | NA | NA | 0.355 | 0.723 | 0.941 | 0.347 | -1.315 | 0.188 |
| **GC:EA** | 64 | NA | NA | -0.423 | 0.673 | 2.001 | **0.045.** | -1.427 | 0.153 |
| **SN:LP** | 64 | NA | NA | -0.416 | 0.667 | 1.919 | 0.055 | -0.912 | 0.362 |
| **GC:LP** | 64 | NA | NA | -1.315 | 0.188 | 1.709 | 0.087 | 0.045 | 0.964 |
| **GC:SN** | 64 | NA | NA | -0.234 | 0.815 | -0.634 | 0.526 | 0.767 | 0.443 |
| **Positively stained cells** | 63 | -1.266 | 0.206 | 0.010 | 0.992 | 0.902 | 0.367 | -0.965 | 0.335 |
| **Low stain intensity** | 63 | 0.713 | 0.482 | 0.225 | 0.800 | -0.496 | 0.624 | 0.020 | 0.984 |
| **Moderate stain intensity** | 63 | -0.853 | 0.394 | -0.497 | 0.619 | 1.056 | 0.291 | -0.629 | 0.529 |
| **High stain intensity** | 63 | -1.533 | 0.125 | 0.204 | 0.839 | 0.737 | 0.461 | -0.453 | 0.651 |

#### FOLLICULAR COUNTS

For an individual there was no significant interaction with *in-utero* sex ratio of that piglet’s litter and their birthweight for the number of recruited follicles (GLMM; t-value=.789, n=34, p=.436). There was a significant interaction in primordial (Gamma GLM; t-value=-2.637, n=34, p=.008) and total (Gamma GLM; t-value=-2.754, n=34, p=.006) follicle numbers per cm^2^ between these two predictor variables i.e. sex ratio *in-utero*. As seen in Figure 6, increasing birth weight was negatively correlated with primordial and total follicle numbers in females from male biased litters. Contrary to this, female-biased and non-biased litters showed no difference in follicle numbers between different weights.

**Figure 6.**
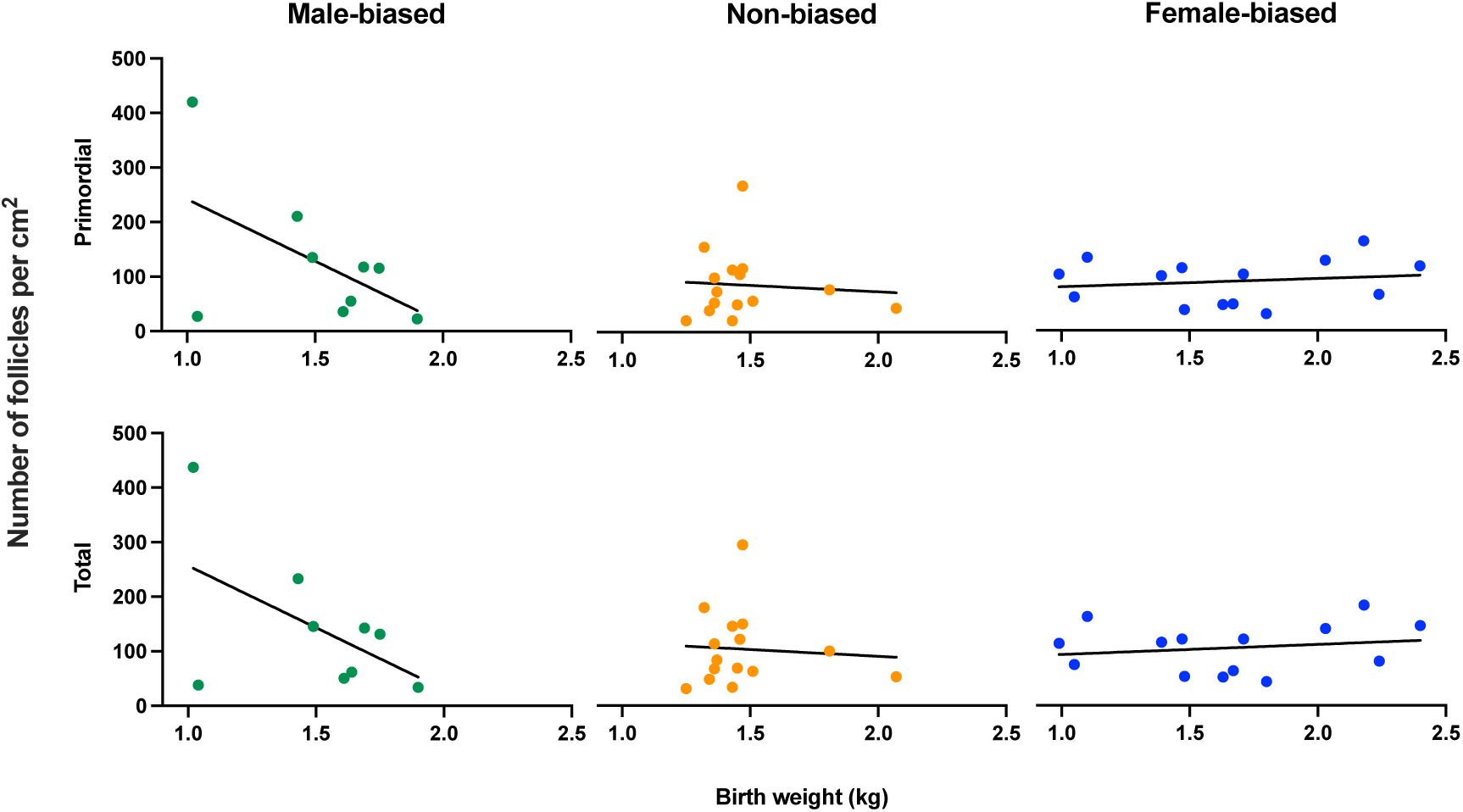
Birth weight of piglets for both primordial and total follicle numbers per cm2. Scatter plots hold fitted regression lines, individual points have been grouped according to the bias of the litter as either female-biased (>65% females), non-biased (45-54.9% females), and male-biased (<35% females), n=34. Pigs from male biased litters held a higher follicle count (both primordial and total) when they held a lower birth weight.

#### UTERINE MORPHOLOGY AND PROLIFERATING CELLS

There was no interaction observed between the sex ratio and birth weight for endometrial area (Gamma GLM; z-value=-1.416, n64, p=.157), nor luminal perimeter (Gamma GLM; z value=-1.595, n=64, p=.1107). No significant interaction was found between birthweight nor sex ratio for the proportion of positively PCNA-stained nuclei (Gamma GLM; z-value=-1.266, n=63, p=.206), nuclei with low stain intensity (GLMM; z-value=.713, n=63, p=.482), nuclei with a moderate stain intensity (Gamma GLM; z-value=-.853, n=63, p=.394), nor the nuclei with a high stain intensity (Gamma GLM; z-value=-1.533, n=63, p=.1252).

### NON-INTERACTIVE EFFECTS OF SEX RATIO, BIRTH WEIGHT, AND SLAUGHTER WEIGHT

#### FOLLICULAR COUNTS

No significant difference in numbers of recruited (GLMM; t-value=.442, n=34, p=.679), nor atretic (Gamma GLM; t-value=.817, n=34, p=.414) follicles were observed between sex ratios. Similarly, both recruited (GLMM; t-value=669, n=34, p=.509) and atretic (Gamma GLM; t-value=1.775, n=34, p=.0759) follicle numbers were not affected by the birthweight of an individual. The slaughter weight had no effect on any aspect of follicle count measured (primordial - Gamma GLM; t-value=860, n=34, p=.589; recruited - GLMM; t-value=.995, n=34, p=.330; atretic - Gamma GLM; t-value=1.468, n=34, p=.142);, nor total - Gamma GLM; t-value=2.754, n=34, p=.611).

#### UTERINE MORPHOLOGY

The sex ratio of the litter from which a pig originated, birth weight, or slaughter weight did not significantly affect the total cross section of endometrial area (Gamma GLM, n=64; z-value=.022, p=.825; z-value=1.376, p=.169; and z-value=.787, p=.431; respectively), size of the luminal area as measured by luminal perimeter (Gamma GLM, n=64; z-value=1.096, p=.273; z-value=1,915, p=.055; z-value=-.151, p=.880; respectively), nor number of stromal cells as measured by nuclear staining (Poisson GLM, n=64: z-value=1.096, p=.2731; z-value=-1.915, p=.0555; z-value=-.151, p=.8799; respectively).

The total endometrial gland numbers (Poisson GLM, n=64; z-value=.117, p=.907; z-value=.058, p=.954; z-value=.054, p=.957; respectively) and larger glands alone (Poisson GLM, n=64; z-value=-1.033, p=.302; z-value=.432, p=.666; z-value=-.908, p=.364; respectively) were not significantly different between individuals of different *in-utero* sex ratios, birth weights, or slaughter weights.

#### PROPORTIONAL ANALYSES

The proportion of the secretory structures were then analysed in proportion to the endometrial area – this was to investigate whether the structures differed between sex ratio litter individuals in regard to the size of their reproductive tracts. The luminal perimeter (cm^2^), glandular count, and stromal nuclei in relation to the endometrial area was analysed and no effects were seen for the ratio of luminal perimeter nor stromal cell number nuclei to endometrial area for females from different sex ratio litters (Gamma GLM, n=64; z-value=.874, p=.382; z-value=.355, p=.723; respectively), birthweight (Gamma GLM, n=64; z-value=-.142, p=.887; z-value=.941, p=.347; respectively), nor slaughter weight (Gamma GLM, n=64; z-value=-1.397, p=.162; z-value=-1.315, p=.188; respectively). However, although the ratio of the total number of glands in the endometrial area for females from different sex ratios (Gamma GLM; z-value=-0.423, n=64, p=.67259), and slaughter weights (Gamma GLM; z-value=-1.427, n=64, p=.15349) were not significantly different, there was a significant difference in effect of the birth weight (Gamma GLM; z-value=2.005, n=64, p=.04492), Figure 7.

**Figure 7.**
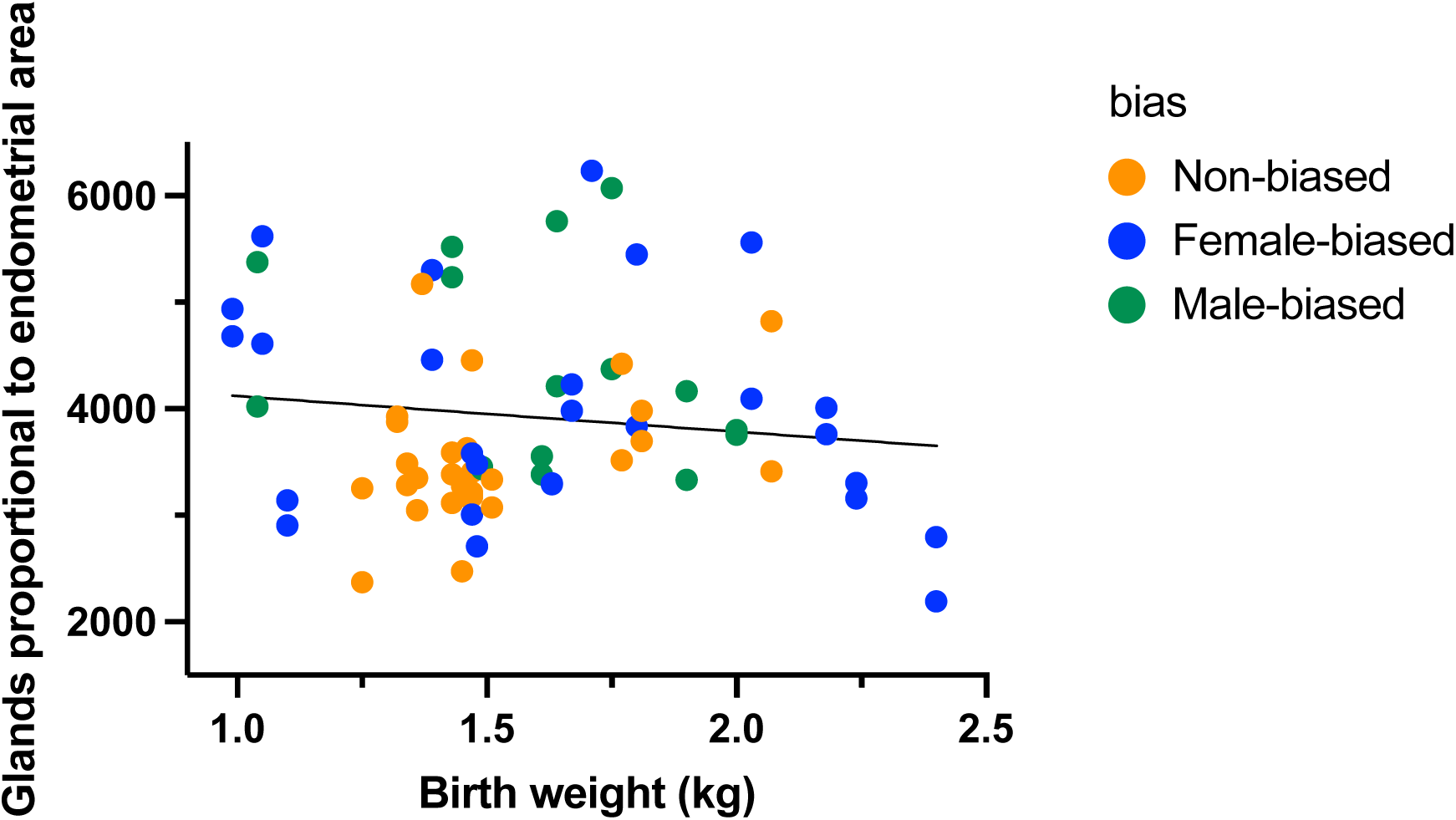
The ratio of glands to endometrial area in female pigs in relation to their birth weight (kg). Ratio given for those individuals gestated in either female-biased (>65% females; Blue dots), non-biased (45-54.9% females; Yellow dots), and male-biased (<35% females; Green dots), n=34 in total.

The analyses investigating the ratio of different secretory tissues against each other within the uterine horn demonstrated no significant differences in ratios between stromal nuclei and luminal perimeter (Gamma GLM, n=64; Sex ratio - z-value=-0.416, p=.6674; Birthweight - z-value=1.919, p=.0550; Slaughter weight - z-value=-.912, p=.3619), gland counts and luminal perimeter (Gamma GLM, n=64; Sex ratio - z-value=-1.315, p=.1884; Birthweight - z-value=1.709, p=.0874; Slaughter weight - z-value=.045, p=.9642), nor gland counts and stromal nuclei (Gamma GLM, n=64; Sex ratio - z-value=-.234, p=.815; Birthweight - z-value=-

.634, p=.526; Slaughter weight - z-value=.767, p=.443).

#### PROLIFERATING CELLS

The sex ratio of the litter from which a pig originated, birth weight and slaughter weight did not show a significant effect on the proportion of PCNA-positive stained cells (Gamma GLM, n=63; z-value=.010, p=.992; z-value=.902, p=.367; z-value=.965, p=.335; respectively), nuclei with a low stain intensity (GLMM, n=63; z-value=.225, p=.800; z-value=-.469, p=.619; z-value=.020, p=.984; respectively), moderate stain intensity (Gamma GLM, n=63; z-value=-.497, p=.619; z-value=1.056, p=.291; z-value=-.629, p=.529; respectively), and high stain intensity (Gamma GLM, n=63; z-value=,204, p=.839; z-value=,737, p=.461; z-value=-.453, p=.651; respectively).

## DISCUSSION

The aim of this study was to investigate how gestation of individuals in a sex biased *in utero* environment altered development and reproductive potential of the female offspring. We hypothesised that gestation of a female in a predominantly male or female in utero environment would alter primordial follicle pool and the development of the uterine capacity.

Irrespective of whether the follicle count was assessed on a per ovary or tissue size basis the outcome was the same. We found that the primordial follicle pool and total follicle count in the ovary was significantly more variable when females originated from a male biased uterine environment. We also found that the follicle counts were affected by the uterine environment differently by those from a male biased vs a female biased uterine environment, based on their birth weight. Previous research in species including mice and pigs suggest an important role for androgens on follicular and CL development. In gilts this is illustrated by increased ovulation rates following dihydrotestosterone treatment [39], and CL dysfunction that is marked by decreased progesterone production when flutamide was used to block androgenic actions [40]. Overall, this study found no effects of sex ratio on the non-ovulatory recruitment, nor atresia of follicles, suggesting that an androgenised uterine environment does not interfere with non-cyclic folliculogenesis, nor the breakdown of follicles in the pre-pubertal commercial pig. Androgens in sheep were found to increase follicular recruitment when offspring were in a hyper-androgenised maternal circulation, resulting in detrimental effects such as multi-follicular ovaries (similar to PCOS) and early cessation of cyclicity exposed to hyper-androgenised maternal circulation in sheep [41]. However, it is important to note that sheep normally only bear 1-3 offspring per gestation and hence aren’t litter bearing. Thereby, they are less likely to be exposed to a biased uterine environment than in the pig, potentially leading to a higher sensitivity to hyper-androgenisation of the uterine environment. This may contribute to the species-specificity observed on androgenisation….

Despite no effects seen in the number of follicles recruited females from male-biased litters had a higher count of primordial and total follicles both per cm^2^ when the individual pig held a low birth weight. However, females with a higher birth weight were found to hold a higher number of primordial and total follicles if from female-biased litters. This suggests that the effect of an androgenised environment and an oestrogenised uterine environment seem to have opposite influences on the development of the primordial follicle pool. This is further affected by the birthweight of the particular individual. An androgenised uterine environment may increase primordial germ cell proliferation, resulting in a larger TOR. A larger TOR would be in accordance with Seyfang and colleagues [42] who found that androgenised female pigs were more likely to ovulate and had higher CL counts when from a male-biased compared to female-biased litters when treated with gonadotrophins at 18 weeks of age. This could be due to in a higher TOR. Research in non-litter bearing species, who will hold different timings of PGC establishment, conflicts with that in pigs; findings in sheep in which individuals from a hyper-androgenised uterine environment displayed lower TOR’s than their control counterparts [41]. This may be due to the pre-mentioned species differences, or due to the study design mimicking maternal testosterone via intramuscular testosterone propionate in pregnant ewes, rather than uterine hyper-androgenisation. This was an unexpected finding, as piglets that are below 1.3 kg at birth have been found to exhibit less competent post-natal development and reduced survivability [43], suggesting a detrimental effect of low birth weight on offspring. Compensatory growth following low birth weights has been found to lead to delayed puberty onset in mice [44]. Low birthweight piglets have also been found to grow less than normal and high birthweight piglets throughout their life course [45], and do not display the “catch-up”, that piglets that have been fed restrictively do [46]. Although our findings suggest a benefit to the TOR in low birthweight piglets, with no effect on the recruitment nor atresia of follicles, there may be long-term implications as indicated in for example mice.

Despite there being, to our knowledge, no previous research specifically investigating the effect of an *in-utero* sex bias on the uterine morphology or efficiency, post-natal uterine gland development is influenced by lactocrine aspects [47] and oestrogenic influences leading to reduced uterine responsiveness to embryotropic signals [30]. However, we did not find any evidence suggesting that gestation of a female in a male-biased uterine environment is detrimental to uterine development in our study. We did not demonstrate any effect of litter sex ratio on any of the defined uterine measures, their ratios, or the cell proliferation. However, it was found that the birth weight of a female pig had a significant effect on the proportion of glands in the uterus, relative to the uterine size. In this instance the higher the birth weight, the lower the proportion of glands. This may be due to a larger uterus rather than specifically lower developed uterine glands. We propose that synchronised cyclic animals should be used to further investigate whether this effect is having a real effect on the reproducing gilts and sows.

The most important outcome that we found in this study is the increased variability in reproductive measures in females originating from male biased litters. In mice, the strain has shown to account for major variation with regard to follicular profiles [48]. The TOR has also been found to be greatly variable within individuals of the same species, with reports of up to 20-fold differences in individuals through the neonatal period and puberty [49], [50]. This would partially clarify the high variation seen in follicular numbers of pigs from male-biased litters in our data, however, it is interesting to note that the variation is considerably smaller in both non- and female-biased litters. There may be an underlying cause of the increased variation of TOR in male biased litters that hold functional consequences not yet understood. What is also evident is that the number of stromal cell nuclei within the endometrium was significantly more variable in females that came from either extreme, male- and female-biased litters. This variability in females from male- and female-biased litters was also seen in the birth weight of pigs. This could potentially impact on stromal-derived growth factors that enhance secretion from the luminal and glandular epithelium [51]. The stromal cells are also key in supporting the underlying implantation structures and hence [51], variability within their numbers could lead to variability in uterine implantation capabilities, and thus survival of the embryo. Variability is known to be a major issue for pig producers for the many reasons and is commonly found in many different aspects of production. Placental efficiency was found to be highly variable in the large white at an approximate level of three-fold [52], with two-fold variation within litters. As indicated in these findings, higher levels of variation was seen in both male-, and female-biased litters, compared to the non-biased litters. This suggests that a biased litter may be a contributor to the variability of reproductive output commonly reported in female pigs. Furthermore, such biased litters were found to lead to more variation in the birth weight of offspring. Selection for larger litter sizes over time has resulted in litters of higher numbers but with low and greatly variable birth weights [53]. Low birthweight piglets from large litters are often cross fostered or euthanized as they will not be able to compete with their larger siblings for teats and have poor pre-weaning survival rates [21].

Under the premise of this study pigs were expected to, at slaughter, not yet have become cyclic. Some pigs may however have reached puberty earlier than the norm, skewing results. Therefore, this suggests that to truly understand the effects of a biased litter on the recruitment and atresia of follicles, future work should utilise synchronised gilts, and these current findings should be cautiously interpreted. There may be an increase in the depletion of TOR in older pigs, which would normally see a 70% decrease from E50 to 300 days after birth [54]. Finally, albeit currently a very novel avenue, there may be an effect of an androgenised uterine environment on the development of oogonial stem cells, if they functionally exist within the pig. Potentially inhibiting a maintenance of the TOR. However, the subjective nature of manual quantification of the TOR must be taken into consideration when interpreting the results from this study. Future research should investigate the TOR of synchronised, pre-pubertal pigs to further understand the effects of a sex biased litter on the follicular profile of commercial pigs. Synchronisation would allow for a detailed understanding of deviating recruitment patterns observed in previous studies, which this experimental design doesn’t hold sensitivity to fully investigate. Further to this study, investigating the depletion of the ovarian reserve over time would help understand the long-term effects that a bias may hold on reproductive longevity.

In conclusion, females that originate from a predominantly male-biased litter i.e. an androgenised uterine environment, and have low birth weight, display increased primordial ovarian reserve along with increased variability of primordial follicle numbers as compared to pigs that originate from non- and female-biased litters. Conversely, a higher birth weight resulted in a greater primordial ovarian reserve if the female pig originated from an oestrogenised uterine environment. Pigs from either litter bias (male or female) were found to have a significantly higher variation in birth weight than if a pig originated from a non-biased litter. These data have implications for reproductive potential of females gestated in sex-biased in utero environments.

## ACKNOWLEDGEMENTS

Microscopy was facilitated by the University of Leeds Bioimaging Facility, and we would also like to thank the National Pig Research Centre for their work in relation to this publication. Work in NF’s group is supported by N8 agri-food pump priming, QR GCRF, UN, LTHT, as well as BBSRC grant number BB/R017522/1.

